# Human oocytes harbouring damaged dna can complete meiosis-i

**DOI:** 10.1101/752113

**Authors:** Gaudeline Rémillard-Labrosse, Nicola Dean, Adélaïde Allais, Aleksandar I. Mihajlović, Shao Guang Jin, Weon-Young Son, Jin-Tae Chung, Melissa Pansera, Sara Henderson, Alina Mahfoudh, Naama Steiner, Kristy Agapitou, Petros Marangos, William Buckett, Jacob Ligeti-Ruiter, Greg FitzHarris

**Author notes:** These authors contributed equally (Gaudeline Rémillard-Labrosse, Nicola Dean).

## Abstract

Chromosomal abnormalities such as aneuploidies and DNA damage are considered a major threat to the establishment of healthy eggs and embryos. Recent landmark studies showed that mouse oocytes with damaged DNA can resume meiosis and undergo Germinal Vesicle Breakdown (GVBD), but then arrest in metaphase of meiosis-I in a process involving Spindle Assembly Checkpoint (SAC) signalling. Such a mechanism could help prevent the generation of metaphase-II (Met-II) eggs with damaged DNA. However we report that this is not the case in the human oocyte. DNA damage prevents human oocytes from undergoing GVBD in some cases. Strikingly however, most oocytes harbouring DNA damage progress through meiosis-I and subsequently extrude the first polar body (PB1) to form a metaphase-II egg, revealing the absence of a DNA-damage-induced SAC response. Analysis of the resulting metaphase-II eggs revealed highly disorganised spindles with misaligned and heavily damaged chromosomes. Our results suggest that DNA damage accumulated in meiosis-I, such as could occur during *in vitro* maturation procedures, does not prevent polar body extrusion and therefore could persist in morphologically normal looking metaphase-II eggs.

## INTRODUCTION

Maintaining genomic fidelity during oogenesis is paramount in the generation of a healthy embryo. Defects to the genome that can threaten reproductive potential include aneuploidies, where cells gain or lose entire chromosomes, and DNA damage where genomic insults cause DNA breakages that can lead to mutation. Both are considered to be hazardous during oocyte maturation, the final stage of oogenesis by which the fully grown germinal vesicle stage oocyte undergoes germinal vesicle breakdown, builds a spindle apparatus, and segregates the chromosomes between the first polar body and the metaphase-II egg. Thus, describing the mechanisms that the oocyte can deploy to sense and deal with aneuploidies and DNA damage during oocyte maturation is key to understanding how healthy eggs are made.

Successful chromosome segregation, and thus avoidance of aneuploidy, is achieved by attaching chromosomes to spindle microtubules via kinetochores, such that chromosomes are pulled correctly in opposite directions during anaphase when the cell divides. The major cellular mechanism that serves to prevent chromosome segregation errors and therefore the generation of aneuploid cells is the Spindle Assembly Checkpoint (SAC). Any unattached kinetochore triggers the formation of an inhibitory signal that prevents anaphase until correct attachment has been attained (Lara-Gonzalez et al., 2012; Musacchio, 2015). However, it has long been realised that the SAC is highly ineffective at preventing aneuploidy in the mammalian oocyte (Greaney et al., 2017; Mihajlović and FitzHarris, 2018). SAC signalling is evidently in operation in the mouse oocyte, since extreme spindle damage can prevent polar body formation, and removal of SAC components shortens meiosis and exacerbates chromosome segregation errors (Homer et al., 2005; McGuinness et al., 2009). However, importantly, interventions that cause moderate spindle defects fail to elicit a SAC-mediated meiosis-I arrest, and oocytes can initiate anaphase chromosome segregation during oocyte maturation despite misaligned chromosomes that could cause aneuploidy (Gui and Homer, 2012; Kolano et al., 2012; Lane et al., 2012; Sebestova et al., 2012). Thus, at least in mouse, the SAC is unusually ineffective at detecting spindle defects in the oocyte (reviewed in (Mihajlović and FitzHarris, 2018)).

Recent experiments examining the impact of experimentally-induced DNA damage upon oocyte maturation led to an unexpected possible explanation for the aforementioned SAC failures in mouse. Multiple groups showed that oocytes harbouring DNA damage readily undergo Germinal Vesicle Breakdown but then arrest in meiosis-I harbouring an apparently normal MI (meiosis I) spindle, thereby failing to extrude the first polar body (Collins and Jones, 2016; Collins et al., 2015; Lane et al., 2017; Marangos and Carroll, 2012; Marangos et al., 2015). These results were unexpected since DNA damage in mammalian somatic cells tends to prevent nuclear envelope breakdown at mitosis entry (equivalent to preventing GVBD), but damage in mitosis does not prevent anaphase onset (Giunta et al., 2010; Rieder and Cole, 1998). Further analysis in mouse oocytes found that the DNA-damage-induced arrest could be over-ridden by inactivation of SAC signalling (Collins et al., 2015; Lane et al., 2017; Marangos and Carroll, 2012). A possible interpretation of this is that the mammalian oocyte has been repurposed to survey DNA damage, rather than the fidelity of chromosome segregation. Such a scenario would hold significant clinical implications, as it implies that the mammalian oocyte possesses a robust mechanism for preventing oocytes harbouring DNA damage incurred during the final stages of oogenesis, such as could occur during *in vitro* maturation or following ovulation induction, from becoming fertilisable metaphase-II eggs. Whether DNA damage prevents completion of oocyte maturation in humans, as it does in mouse, is as yet unknown.

## MATERIALS AND METHODS

### Oocyte collection and culture

Mouse oocytes were collected at the germinal vesicle stage from the ovaries of CD1 females (Charles River) aged ∼3months, 44-48 hours after intra-peritoneal injection of 5IU of pregnant mares serum gonadotrophin (PMSG, GENWAY BIOTECH), as previously (Haverfield et al., 2017; Nakagawa and FitzHarris, 2017). Oocytes were released into M2 media containing 200μM of 3-isobutyl-I-methylxanthine (IBMX, Sigma). Oocyte culture was performed in M16 medium covered in mineral oil, and oocyte maturation triggered by washing oocytes through a minimum of 5 drops of IBMX-free M16. Oocyte culture was performed at 37°C in an environment of 5% CO_2_ balanced with air.

Human oocytes were collected from women undergoing IVF treatment with intra-cytoplasmic sperm injection. 50 patients aged from 27-44 years old (Average = 37 years old) consented to the study at the MUHC Reproductive Center (888 Boul de Maisonneuve E #200, Montréal, QC H2L 4S8) donating a total of 149 oocytes at the germinal vesicle (GV) or meiosis I (MI) stage. Human oocytes were cultured in Global total media for fertilization (Life Global Group) supplemented with 0.75 IU/ml of menotropin (Repronex) covered in mineral oil, in an incubator with 6% CO_2_ at 37°C.

### Chemical treatments

Media was supplemented with the following chemicals as described in the Results section: etoposide (100 µg/ml in DMSO; Sigma Aldrich), neocarzinostatin (0-5 µg/ml in MES buffer; Sigma Aldrich), monastrol (200 µM in DMSO; Calbiochem).

### Image acquisition

Mouse oocytes were fixed using 2% paraformaldehyde/PBS for 20 min and permeabilized for 10 minutes with 0.25% Triton-X/PBS, at room temperature. Human oocytes were fixed using 2% paraformaldehyde + 0.05% triton-X/PHEM for 15 min and permeabilized for 15 minutes with 0.05% Triton-X/PHEM, at room temperature. Primary antibodies used as follows: α-tubulin (Sigma, 1:1000), γH2AX (Trevigen, 1:800). Alexa-labeled secondary antibodies (Life Technologies, 1:1000) were used for detection. DNA was labeled using Hoechst (Hoechst 33342, Sigma, 1:1000) and actin was detected using Alexa647-phalloidin or Alexa555-phalloidin (Invitrogen, 1:300). Fixed cells imaging was performed on a Leica SP8 confocal microscope fitted with a 20x 0.75NA objective and a HyD detector. Image acquisition was performed via the LASX software. Live cell imaging to measure GVBD and PBE timing was achieved on a Zeiss Axio observer, with an axiocam and 5X objective using bright field imaging. Image acquisition was performed via the ZEN pro software. Where possible, control and drug-treated groups were imaged side-by-side with the same pre-imaging manipulations and imaging settings.

### Analysis and Statistics

All data analysis was performed using ImageJ/Fiji. Fluorescence intensity measurements for analysis of γH2AX staining was performed in ImageJ. Total fluorescence within the nucleus/spindle region was integrated in 3 dimensions, and background subtraction performed using a neighbouring equal-sized (DNA-free) cytoplasmic region. Where shown, error bars represent SEM. All data analysis was performed using Excel and/or GraphPad Prism, statistical tests used are accordingly noted in figures legends.

### Ethics approval

Mouse experiments were approved by the Comité Institutionel de Protection des Animaux du CRCHUM (CIPA; project IP18037GFs). Handling of all human material was approved by the Comité d’éthique de la Recherche du CHUM (Project 14.157). All patients consented to the donation of their GV oocytes to the study.

## RESULTS

### Impact of etoposide upon mouse and human oocytes

Prior to examining the impact of DNA damage upon human oocytes, we first sought to confirm that the previously reported effects of DNA damage upon mouse oocyte maturation were observed in our laboratory. We first employed upon etoposide, a topoisomerase-II inhibitor that induces DNA double strand breaks and has been shown by multiple groups to prevent first polar /body extrusion in mouse (Collins et al., 2015; Marangos et al., 2015). As previously reported, a 1 hour exposure of IBMX-treated GV-arrested mouse oocytes to etoposide caused a dramatic increase in nuclear γH2AX immunoreactivity, indicative of DNA damage (Figure 1A,B). To assess the impact of the damage on oocyte maturation, oocytes were treated with etoposide either acutely prior to removal of IBMX, or throughout oocyte maturation. As previously reported, etoposide slowed but did not potently prevent GVBD, but more potently affected polar body extrusion (Collins et al., 2015; Marangos and Carroll, 2012; Marangos et al., 2015). Whereas an etoposide pre-incubation slowed polar body extrusion, continuous exposure robustly prevented PB1 formation (Figure 1C,D). Thus, in broad agreement with other groups, etoposide does not prevent GVBD, but inhibits polar body formation in mouse.

**Figure 1:**
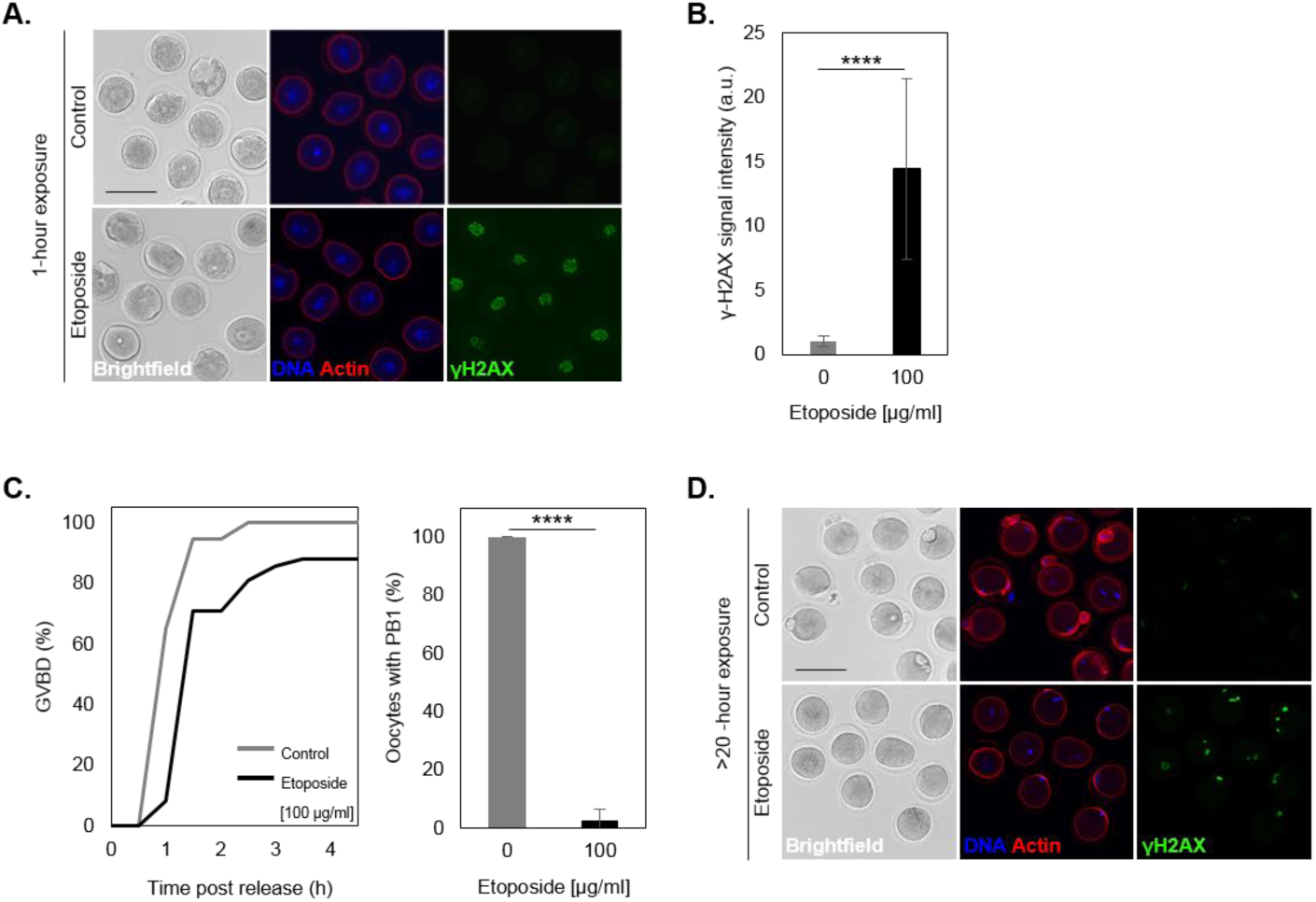
Etoposide triggers an arrest in MI in mouse oocytes. **A.** Mouse oocytes collected at GV stage were incubated with DMSO only (control) or 100 µg/ml of etoposide. Cells were then fixed after 1 hour +/- etoposide. Representative images displaying bright field, DNA (Hoechst), actin (Alexa-phalloidin) and DNA damage (γ-H2AX antibody) are shown. Bars represent 100 µm. **B.** DNA damage was assessed via measurement of the γ-H2AX signal intensity in each cells incubated +/- drug for 1 hour. Data comprises 19 control and 24 etoposide treated oocytes. **C.** Germinal vesicle breakdown (GVBD, left panel) and polar body extrusion (PB1, right panel) were evaluated for both groups (control and etoposide) at different times after IBMX release. Data comprises 37 control and 40 etoposide-treated oocytes. Error bars represent SEM. P-values were obtained via an unpaired Student’s t-test and **** indicates p<0.0001. **D.** Mouse oocytes collected at GV stage were incubated with DMSO only (control) or 100 µg/ml of etoposide. Cells were then fixed after 20 hours of maturation +/- etoposide. Representative images displaying bright field, DNA (Hoechst), actin (Alexa-phalloidin) and DNA damage (γ-H2AX antibody) are shown. Bars represent 100 µm.

To determine whether DNA damage induced by etoposide could prevent oocyte maturation in humans we used GV stage oocytes recovered for routine ICSI cycles and donated to research. We previously reported that ∼70% of oocytes recovered in this manner extrude a polar body to become morphologically-normal metaphase-II eggs 24-48 hours after collection in our standard incubator culture conditions (Haverfield et al., 2017). We first confirmed that a 1 hour incubation of (100μg/ml) etoposide induced DNA-damage using γH2AX antibodies. As expected, fluorescence quantification confirmed substantial DNA-damage within 1 hour of etoposide treatment (Figure 2A,B). Next we set out to determine the effect of etoposide treatment on oocyte maturation. Therefore, in order to ensure high levels of DNA damage throughout oocyte maturation we incubated oocytes in etoposide continuously, similar to mouse oocytes. As previously (Haverfield et al., 2017), ∼70% of control (DMSO treated) oocytes extruded polar bodies within 48h of oocytes collection, most of those within the first 24h. Strikingly, a statistically significant proportion of etoposide-treated oocytes remained arrested at GV stage (40%, P<0.05). Most notably, however, of those etoposide-treated oocytes that underwent GVBD, almost all extruded a polar body (Figure 2C). Confocal imaging confirmed that the chromatin in those etoposide-treated oocytes that extruded a polar body was indeed substantially damaged as judged by intense γH2AX labelling, compared to contemporaneous vehicle controls (Figure 2D). No prevention of polar body formation was also observed in a separate small series of experiments where etoposide was applied following GVBD (Figure 2D bottow row). Thus etoposide induces DNA damage in human oocytes, preventing a subset from undergoing GVBD, but the majority of etoposide-treated oocytes progress to metaphase-II despite substantial DNA damage.

**Figure 2:**
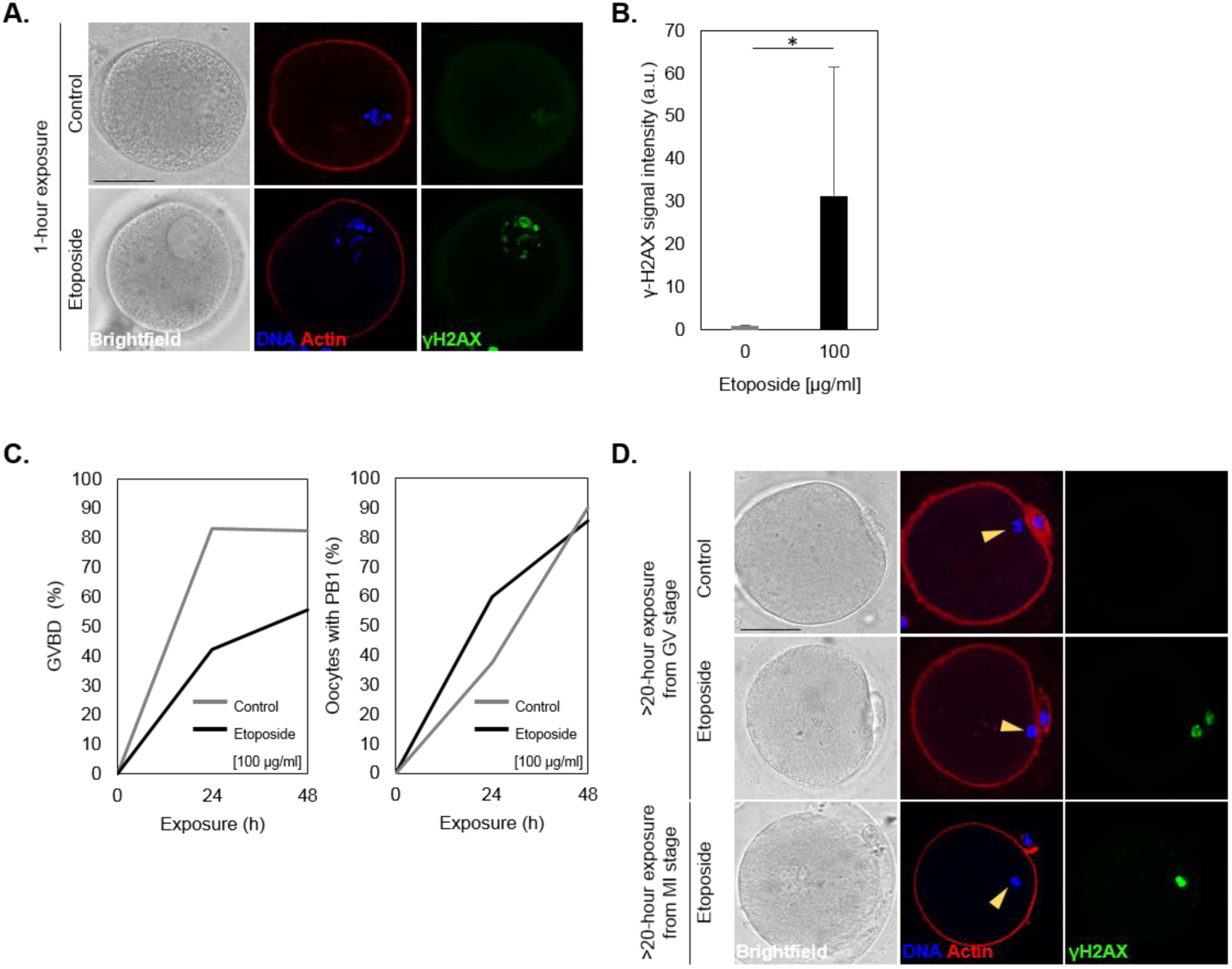
Etoposide does not arrest human oocytes in MI, despite causing DNA damage. **A.** Human oocytes collected at GV stage were incubated with DMSO only (control) or 100 µg/ml of etoposide. Cells were then fixed after 1 hour +/- etoposide. Representative images displaying bright field, DNA (Hoechst), actin (Alexa-phalloidin) and DNA damage (γ-H2AX antibody) are shown. Bars in images represent 50 µm. **B.** DNA damage was assessed via measurement of the γ-H2AX signal intensity in each cells incubated +/- drug for 1 hour. Data comprises 6 control and 9 etoposide-treated oocytes. Error bars represent SEM. * indicates P<0.05 (Students Ttest). **C.** Germinal vesicle breakdown (GVBD, left panel) and polar body extrusion (PB1, right panel) were evaluated for each oocyte from both groups (control and etoposide) at different time points after addition of drug (or DMSO) at the GV stage. For control group (n)=18 oocytes and for etoposide treated group (n)=22 oocytes from 14 patients. Note that percentage of PB1 extrusion is expressed as a proportion of oocytes that underwent GVBD. **D.** Human oocytes collected at GV stage were incubated with DMSO only (control) or 100 µg/ml of etoposide. Cells were then fixed after 24-48 hours of maturation +/- etoposide. Note that the control and treated oocyte come from the same patient and were processed and imaged contemporaneously with identical microscope settings, and that there is a high level of DNA damage in etoposide-treated oocytes. Note that the lower panel in D shows a separate experient in which oocytes were incubated with etoposide once they reached MI stage. Representative images displaying bright field, DNA (Hoechst), actin (Alexa-phalloidin) and DNA damage (γ-H2AX antibody) are shown. Bars in images represent 50 µm. Yellow arrowhead indicates cellular DNA.

### Impact of neocarzinostatin upon mouse and human oocytes

Next we set out to determine whether a second mechanistically distinct means of introducing DNA damage might prevent polar body extrusion, employing the radiomimetic agent neocarzinostatin (NCS) that has been employed in mouse oocytes previously (Mayer et al., 2016; Yuen et al., 2012). In mouse oocytes, as previously reported (Mayer et al., 2016; Yuen et al., 2012), a 1 hour NCS exposure induced readily detectable γH2AX signals in GV stage oocytes, similar to etoposide (Figure 3A,B). As with etoposide, NCS failed to prevent GVBD, but impacted PB1 extrusion. Consistent with previous studies (Mayer et al., 2016; Yuen et al., 2012) the inhibition of PB1 extrusion was far less robust than with etoposide, resulting in a delay rather than long term inhibition, the majority of NCS-treated oocytes eventually extruding a polar by 12h after release from IBMX (Figure 3C,D). In human oocytes, similar to mouse oocytes, a 1 hour exposure to NCS induced clear γH2AX signals in the human oocyte GV (Figure 4A,B). Oocytes matured in NCS exhibited a minor (non-significant) delay in GVBD timing that was less pronounced than with etoposide. Strikingly though, similar to etoposide, the vast majority of NCS treated oocytes extruded a polar body within 48 hours (Figure 4C). Importantly, analysis of the resulting Met-II oocytes revealed pronounced γH2AX immunofluorescence (Figure 4D). Thus NCS-treated oocytes readily extrude PB1 despite pronounced DNA damage.

**Figure 3:**
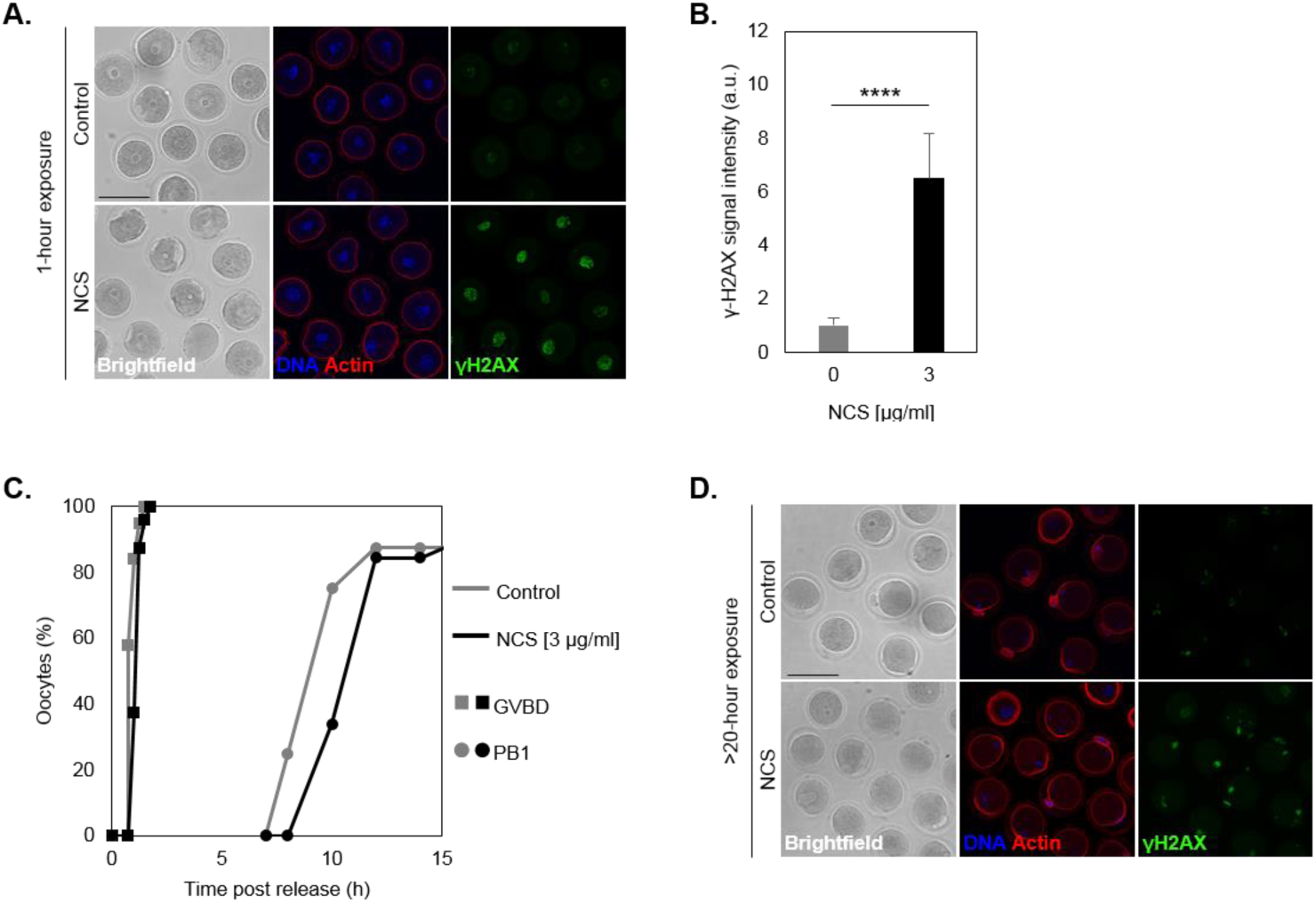
Neocarzinostatin perturbs mouse oocytes maturation. **A.** Mouse oocytes collected at GV stage were incubated with MES buffer only (control) or 3 µg/ml of NCS. Cells were then fixed after 1 hour +/- NCS. Representative images displaying brightfield, DNA (Hoechst), actin (phalloidin) and DNA damage (γ-H2AX antibody) are shown. Bars represent 100 µm. **B.** DNA damage was assessed via measurement of the γ-H2AX signal intensity in each cells incubated +/- drug for 1 hour. Data comprises 15 control and 21 NCS-treated oocytes. Error bars represent SEM. P-value was obtained via an unpaired Student’s t-test and **** indicates p<0.0001. **C.** Germinal vesicle breakdown (GVBD) and polar body extrusion (PB1) were evaluated for each oocyte from both groups (control and NCS) at different time points after IBMX release. **D.** Mouse oocytes collected at GV stage were incubated with MES buffer only (control) or 3 µg/ml of NCS. Cells were then fixed after 24 hours of maturation +/- NCS. Representative images displaying bright field, DNA (Hoechst), actin (phalloidin) and DNA damage (γ-H2AX antibody) are shown. Bars represent 100 µm.

**Figure 4:**
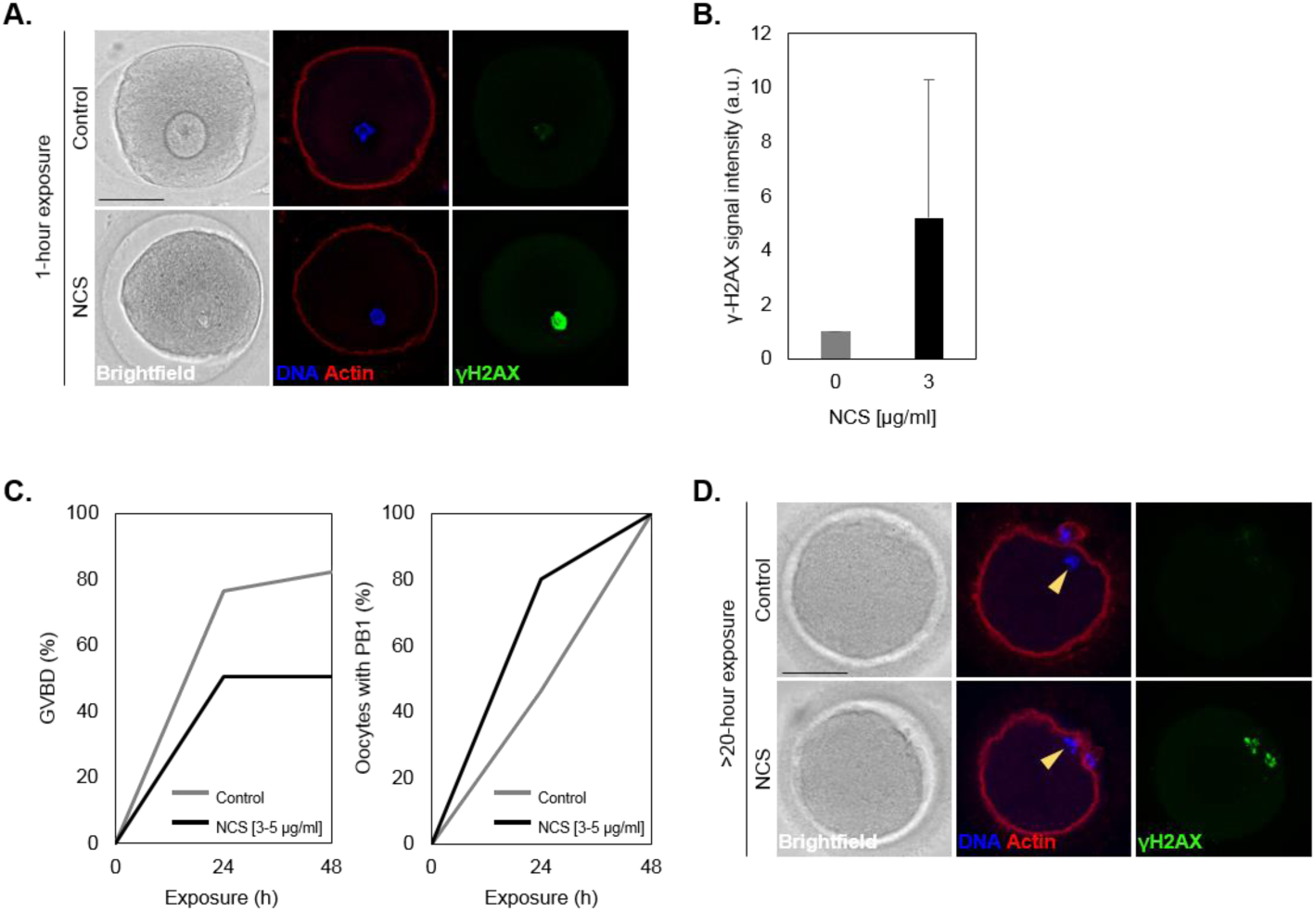
Neocarzinostatin does not arrest human oocytes in MI. **A.** Human GV oocytes were incubated with MES only (Control) or 3-5 µg/ml of NCS. Cells were then fixed after 1 hour of maturation +/- NCS. Representative images displaying bright field, DNA (Hoechst), actin (Alexa-phalloidin) and DNA damage (γ-H2AX antibody) are shown. Bars in images represent 50 µm. **B.** γ-H2AX signal intensity was measured in each cells fixed after 1 hour. Data comprises 4 control and 6 NCS-treated oocytes. Error bars represent SEM. **C.** Oocytes undergoing germinal vesicle breakdown (GVBD, left panel) and polar body extrusion (PB1, right panel) were scored for both groups (Control and NCS). Data comprises 15 control and 19 NCS treated oocytes. Note that percentage of PB1 extrusion is expressed as a proportion of oocytes that underwent GVBD. **D.** Human GV oocytes were incubated with MES only (Control) or 3-5 µg/ml of NCS. Cells were then fixed after 24-48 hours of maturation +/- NCS. Representative images displaying bright field, DNA (Hoechst), actin (Alexa-phalloidin) and DNA damage (γ-H2AX antibody) are shown. Bars in images represent 50 µm. Yellow arrowhead indicates cellular DNA. Note that the control and treated oocyte come from the same patient and were processed and imaged contemporaneously with identical settings.

### Morphology of spindles in DNA-damaged human metaphase-II eggs

Next we sought to determine the integrity of the chromosomes and spindle apparatus in oocytes that had been exposed to etoposide or NCS during oocyte maturation. Chromosomes and spindles were labeled with Hoechst and tubulin antibodies, respectively, and the cell boundary labeled with Alexa-Phalloidin such that oocyte chromosomes could be distinguished from the polar body, and γH2AX antibodies used to observe DNA damage (Figure 5). Vehicle-treated controls that we were able to examine at Met-II all had bipolar metaphase-II spindles. Of these, half (4/8) had fully aligned chromosomes, the other half having varying degrees of chromosome misalignment from the metaphase plate. Etoposide and NCS-treated oocytes had more severe spindle phenotypes. Six out of 11 treated ooctres examined had huighly disorganised spindleas, and only three out of eleven had normal spindles with aligned chromosomes. In some cases chromosomes appeared unusually clustered as a ball. The appearance of chromosomes was qualitatively different depending upon the treatment, with chromosomes in NCS-treated oocytes having a more granular appearance, perhaps suggesting more frequent chromosome breakage. Thus, DNA damage inflicted during oocyte maturation by etoposide or NCS does not prevent polar body formation, but the resulting metaphase-II oocytes harbour damaged chromosomes and frequently defective Met-II spindles.

**Figure 5:**
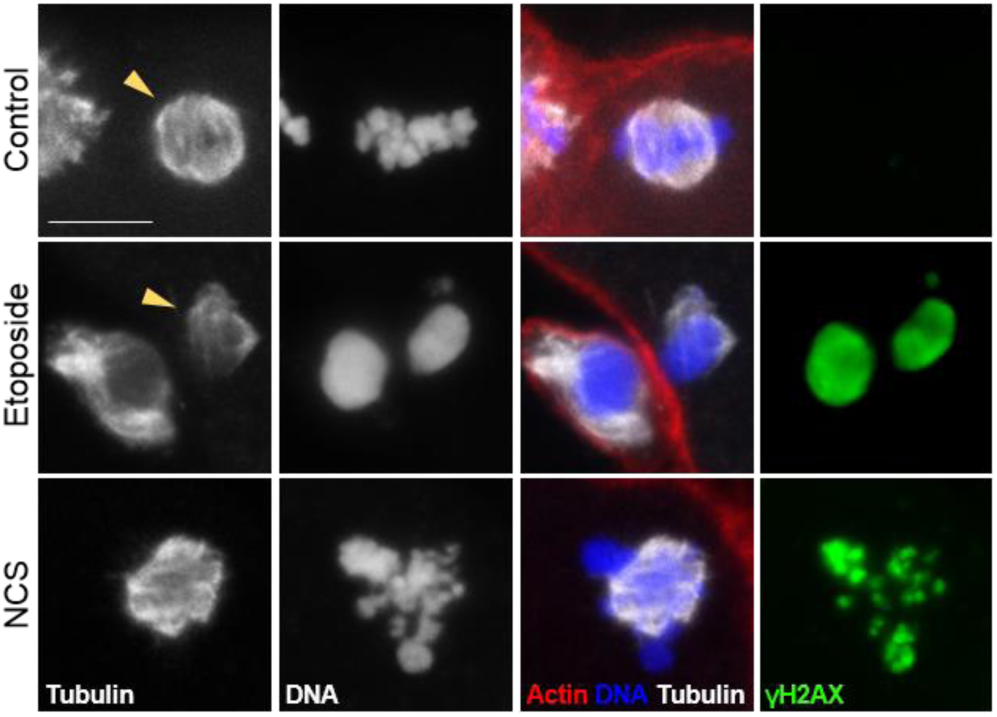
NCS and Etoposide affect metaphase II spindle morphology. Oocytes were incubated with vehicle only (Control) or etoposide [100 µg/ml] or NCS [3 µg/ml]. Cells that matured to MII phase were stained for tubulin (white), DNA (blue), actin (red) and DNA damage (green). Data comprises 8 control, 4 etoposide- and 7 NCS-treated oocytes. See text for further details. Bars represent 10 µm. Yellow arrowhead indicates intracellular tubulin.

### Pharmacological spindle disruption prevents polar body extrusion

In mouse oocytes the SAC can be activated and polar body extrusion prevented by two types of specific intervention; either by DNA damaging agents (Collins and Jones, 2016), or by extreme spindle disruption (Collins and Jones, 2016; Duncan et al., 2009). Our data demonstrates that DNA damage does not robustly activate SAC. Therefore, we set out to determine whether loss of spindle organisation might prevent polar body extrusion. The kinesin-5 inhibitor monastrol caused spindle collapse and the formation of a monopolar spindle in meiosis-I (Figure 6), similar to mouse oocytes (Fitzharris, 2009; Schuh and Ellenberg, 2007). Importantly, whereas 70% of DMSO-treated controls progressed to MetII accrooss this and our previous study, almost all monastrol-treated oocytes remained arrested at MI 48 hours after oocyte collection. Thus severe spindle disruption prevents polar body extrusion, suggesting that Spindle Assembly Checkpoint can, at least in extreme circumstances, be activated in human oocytes sufficiently to prevent polar body extrusion.

**Figure 6:**
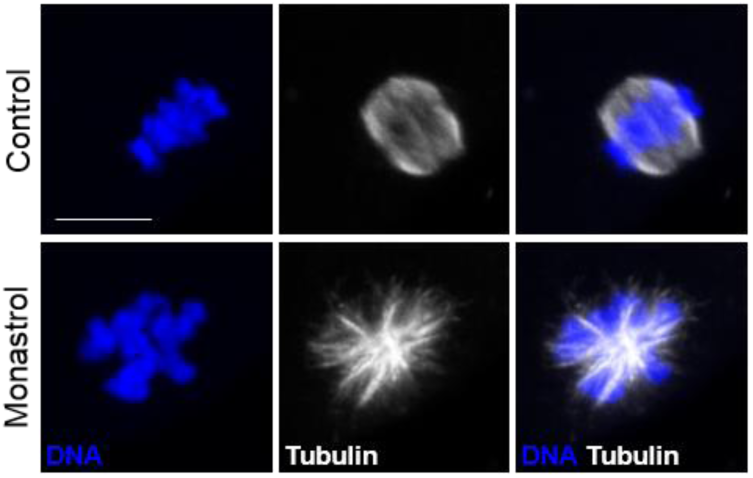
Monastrol affects spindle morphology and blocks polar body extrusion in human oocytes. Human MI-stage oocytes were incubated with DMSO only (Control) or monastrol [200 µM]. Cells were then fixed after 24-48 hours and stained for DNA (blue) and tubulin (white). Bars represent 10 µm. Note that polar bodies were seen in only 2 out of 9 monastrol-treated oocytes, compared with 70% of PB1 extrusion seen in control oocytes throughout this and our previous studies (Haverfield et al., 2017).

## DISCUSSION

The function of the spindle checkpoint in mammalian oocytes has long been a source of contention. In mouse, whereas severe spindle disruption activates the checkpoint to prevent polar body extrusion, moderate spindle insults fail to prevent polar body extrusion, causing defective chromosome segregation that likely leads to aneuploidy (discussed in (Greaney et al., 2017; Mihajlovic and FitzHarris, 2018)). But little has been known about the spindle checkpoint in human oocytes. Live imaging from us and others showed that misaligned chromosomes and spindle defects are common in human oocytes at the time of anaphase onset (Haverfield et al., 2017; Zielinska et al., 2015), indicating that, if it is present at all, SAC is ineffective at preventing chromosome segregation errors, as in mice. Other experiments in human oocytes showed that the SAC component BUB1 can localise to kinetochores in a manner dependent upon MPS1 kinase, alluding to the presence of SAC signalling machinery (Lagirand-Cantaloube et al., 2017). Our monastrol data (Figure 6) provides a simple but clear indication that severe loss of spindle organisation can activate SAC and prevent polar body extrusion. Therefore taken together with the aforementioned work, it would appear that in human oocytes as in mouse, aspects of the pathway are functional, but SAC is insensitive to prevent segregation errors, providing at least part of the explanation for the high levels of aneuploidy in the human oocyte (Gui and Homer, 2012; Hassold and Hunt, 2001; Mihajlovic and FitzHarris, 2018).

Recently, however, it was shown that DNA damage, which does not typically activate SAC in other cell types, apparently does so in mouse oocytes, suggesting the checkpoint has been adapted in female germ-cells to prevent DNA damage rather than aneuploidy (Collins and Jones, 2016). Our main finding here is that the same principle does not appear to apply in human oocytes. Two different DNA damage-inducing agents failed to prevent polar body extrusion despite chronic exposure and clear γH2AX-indicated DNA damage. Simultaneous to the main experimental investigation carried out in Montreal, Canada, similar experiments were performed in Ioanna, Greece. Importantly, parrallel results were obtained in that separately conducted study, high concentrations of etoposide causing substantial DNA damage without preventing polar body formation. Thus similar observations have been made in two different labs (Figure S1). The reason for the distinct outcomes in mouse and human are unclear, as is a molecular explanation as to how DNA damage activates SAC in mouse. A more detailed understanding of the pathway in mouse may facilitate a better understanding of the apparent species difference in oocyte DNA response.

With the number of clinical cycles involving *in vitro* oocyte maturation increasing (Shirasawa and Terada, 2017), understanding the causes and consequences of DNA damage in oocyte maturation is pertinent.

Multiple aspects of the clinical IVF environment could potentially damage the DNA of cultured oocytes and embryos including sub-optimal culture that could cause free radical generation (Burton et al., 2003; Mantikou et al., 2013), media contamination (Kastrop et al., 2007), and exposure to damaging wavelengths of light (Daniel, 1964; Takenaka et al., 2007). Our study of human oocytes employed two potent DNA damage agents applied chronically at high concentrations revealing that even oocytes harbouring DNA damage far exceeding anything that would be expected in any clinical environment managed to extrude a polar body. This interventional approach illustrates however that, in contrast to mouse, even likely very high levels of damage are unable to prevent polar body formation in human oocytes. Thus the landmark observations in mouse should not be extrapolated and interpreted to mean that DNA-damaged oocytes are naturally selected against in the clinic.

## AUTHOR’S ROLES

GF conceived the study, obtained ethical approval, and wrote the manuscript. GF, GRL, ND, AA, AM, KA, PM collected and analysed data. GF, GRL, WB, ND, AA, AM, PM, JLR revised the manuscript. AM, JTC, MP, SH, SGJ, WYS prepared human oocytes for the study. JLR, NS, WB informed and recruited patients. WB is the medical director of the clinic where patients were recruited.

## FIGURES

**Supplementary Figure 1:**
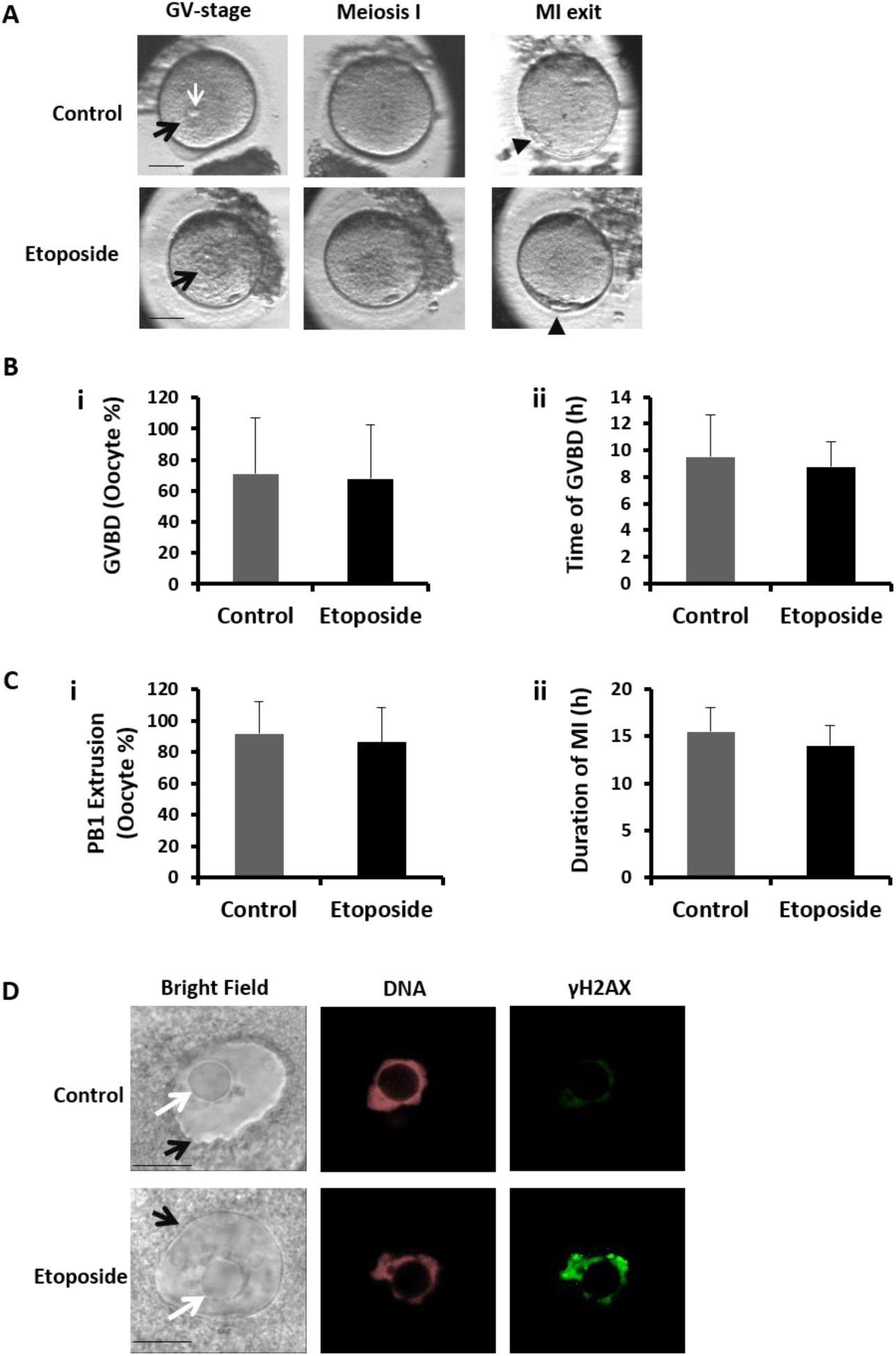
Acute exposure to Etoposide does not activate DNA damage checkpoints in Meiosis I of human oocytes. Germinal Vesicle (GV)-stage human oocytes were treated with 20 μg/ml etoposide for 1h. Treated and control oocytes were placed in the incubator at 37.5°C under 7.1% CO_2_ and cultured in Global for Fertilization + LGPS medium (Life Global Group). Time-lapse monitoring began immediately after oocyte transfer to the incubator, 5h after oocyte recovery by the use of the Primo Vision (TK Biotech) time-lapse culture system. Images were taken every 10 minutes. **A.** Representative images of control and etoposide-treated oocytes prior to oocyte maturation (GV-stage), after GVBD (MI) and after PB1 extrusion (MII). Bars in images represent 50 μm. **B.** The proportion (i) and timing (ii) of GVBD is not affected by the induction of DNA damage by the use of etoposide. 8 experiments involving 18 control and 21 etoposide-treated oocytes, from 8 individual donors were performed in *i* and 6 experiments involving 12 control and 12 etoposide-treated oocytes from 6 individual donors were performed in *ii.* The time of GVBD is measured from the moment of oocyte recovery from the follicle. **C.** The proportion of PB1 extrusion (i) and duration of MI (ii) (from GVBD to PB1 extrusion) is not affected by the use of etoposide. 6 experiments involving 13 control and 14 etoposide-treated oocytes from 6 individual donors were performed in *i* and 6 experiments involving 11 control and 10 etoposide-treated oocytes from 6 individual donors were performed in *ii*. There was no statistical significance in GVBD or PB1 extrusion as a result of etoposide treatment. **D.** Treatment with 20 μg/ml etoposide for 1h causes DNA damage in human oocytes. γH2AX is used as an immunofluorescence marker for the detection of double-strand breaks. Draq7 is used for DNA staining. GV-stage oocytes were fixed immediately after treatment with or without etoposide. Representative confocal images are shown. Bars in images in *D* represent 20 μm. Black arrows: GV; white arrows: nucleolus; arrowhead: (PB1). Human oocytes were recovered and identified as GV-stage oocytes from patients undergoing ICSI cycles. This work was performed with the approval (443/31.7.2018) of the Greek National Authority of Medical Assisted Reproduction.

